# Targeting intracellular mycobacteria using novel antibiotic-loaded nanoparticles

**DOI:** 10.64898/2026.05.14.725169

**Authors:** Pooja Agarwal, Hannah Burnage, Robert Dallmann, Sébastien Perrier, Meera Unnikrishnan

**Affiliations:** Warwick Medical School, University of Warwick, Coventry CV4 7AL, UK; Department of Chemistry, University of Warwick, Coventry CV4 7AL, UK; Faculty of Pharmacy and Pharmaceutical Sciences, Monash University, Parkville, VIC 3052, Australia

**Keywords:** tuberculosis, pH responsive, mannose, macrophages, nanoparticles, rifampicin

## Abstract

Tuberculosis (TB), caused by *Mycobacterium tuberculosis* (*Mtb*), remains a significant global health challenge. Currently treatment of drug-sensitive TB, involves a six-month regimen consisting of a combination of four anti-TB drugs, with drug-resistant TB requiring over two years of treatment and additional drugs. As toxicity of anti-TB drugs often leads to poor compliance, disease relapse and the emergence of drug-resistant strains, new strategies to reduce drug toxicity and shorten treatment duration are critical. We report nanocarrier-based drug delivery systems targeting macrophages, which primarily support replication and survival of *Mtb*. We have developed mannose-functionalized nanoparticles that bind to mannose receptors on macrophages and feature a pH-sensitive core which releases an encapsulated drug in the acidic lysosomal environment of macrophages. Rifampicin (RIF), a main anti-TB drug currently in use clinically, was encapsulated within the nanoparticles. We demonstrate that antibiotic-containing nanocarriers efficiently accumulated in macrophages without causing toxicity. Encapsulated RIF showed enhanced efficacy against both BCG and *Mtb* in primary macrophages. Biodistribution studies in mice revealed that the nanoparticles have extended circulation time and do not induce toxicity. In addition, the encapsulated RIF showed better targeting of mycobacteria when compared to free RIF in a murine model of mycobacterial infection. Such an enhanced bacterial killing using mannose-functionalised nanocarriers loaded with the key anti-TB drug rifampicin offers excellent potential for TB therapy.

## Introduction

*Mycobacterium tuberculosis (Mtb)* is the main causative agent of tuberculosis (TB), which still stands out as one of most deadly diseases worldwide, which is world’s leading cause of death from a single infectious agent. The World Health Organization (WHO) reported that in 2024, approximately 10.7 million people developed tuberculosis (TB), with 1.23 million deaths attributed to the disease^1^ and 390,000 people developing multidrug-resistant or rifampicin-resistant TB (MDR/RR-TB)^1^. TB accounts for higher mortality especially in developing countries, mainly due to malnutrition, low immunity and HIV coinfection^2,3^

*Mtb* is spread in the form of aerosol droplets released by an infected person typically through coughing,and inhaled by a healthy person into their lungs. *Mtb* is taken up by alveolar macrophages in the lungs, which recruit other immune cells eventually resulting in lesion known as granuloma, a hallmark of TB^3,4^. Primary TB is treated by a combinational therapy for six months by first line treatment drugs which includes rifampicin (RIF), isoniazid (INH), ethambutol (ETB), pyrazinamide (PZA) for two months followed by RIF and INH alone for the remaining four months^5^. These antibiotics exhibit toxic effects leading to poor compliance and adherence to the therapy by the patients, which may then increase the risk of development of drug-resistant TB^6^. Multidrug-resistant (MDR) TB is incurable by first-line treatment and is treated by second line drugs; while extensively drug-resistant (XDR) TB is resistant to fluoroquinolones and second-line drugs, with treatment typically requiring therapy with highly toxic drugs^7^.

For successful treatment, TB drugs need to efficiently infiltrate the complex TB-lesions and cross the cell membrane of macrophages, and poor cell permeation of antibiotics is one of the reasons for lengthy treatment duration^5,8,9^. Moreover, subinhibitory intracellular antibiotic concentrations have also been reported to promote drug resistance by upregulating drug-export pumps in *Mtb*^10–12^. Therefore, to reduce treatment time and to prevent development of antibiotic resistance, developing new strategies for more effective delivery of drugs is essential^5^.

Nanomedicine, which utilises nanoparticles for drug delivery, is considered an emerging method to alleviate the problem associated with standard TB therapy^13^, by improving permeability of drugs, reducing drug toxicity, and increasing bioavailability by precise targeting. Drugs are generally conjugated or encapsulated in the core of nanoparticles, but can also be attached to their surface^14^. An exciting feature of this approach is the versatility of the nanocarrier which can be customised to the therapy requirement, e.g. functionalisation with a ligand specific to a target receptor for cell specific drug delivery, or particle core engineered to be pH responsive so that drugs are released at local pH (i.e. acidic lysosomal pH). This approach has been successfully used extensively, for instance in the encapsulation of doxorubicin for the treatment of cancer^15^, or in cystic-fibrosis patients, where nanoparticles have exhibited better diffusion of tobramycin through the thick mucus layer in lungs^16^. Relevant to this work, nanoparticles (either polymeric or lipidic) have demonstrated targeted, pH-activated release of anti-TB drugs with improved killing efficacy *ex vivo* and *in vivo*^17,18^. In these studies, anti-TB drugs RIF and INH stay in blood plasma up to 48 hours, a large increase when compared to free drugs, which get cleared within 24 hours^19^. It is therefore clear that using nanocarrier methodology, the bioavailability of drugs is enhanced, thus offering the potential to reduce their dose and toxicity, and intracellular accumulation, thereby increasing efficacy^20,21^.

In this study we synthesise and characterise dual-targeting nanoparticles which recognise macrophages through specific receptors and release drug upon encounter of lower pH inside the cell to avoid drug release before reaching the target. Using this dual targeting approach, we compared efficacy of nanoparticles encapsulated with rifampicin (RIF), to free drug RIF in killing *Mtb* within macrophages *ex vivo* and in murine infection models, demonstrating enhanced killing of RIF-encapsulated nanoparticles.

## Materials and Methods

### Polymers

#### Materials

(2,2’-Azobis(2-methylpropionitrile (AIBN, 98%), butyl methacrylate (BMA, 99%), 2(Diisopropylamino)ethyl methacrylate (DPAEMA, 97%), 1,2,3,4,6-Penta-O-acetyl-α-D-mannopyranose were purchased from Sigma Aldrich. All monomers were passed through basic aluminium oxide before used to remove inhibitors. N-hydroxy succinimide (NHS, 98%), 1-ethyl3-(3’ dimethylaminopropyl)carbodiimide hydrochloride (EDC.HCl), sodium methoxide (NaOMe, 5.4 M, 30 wt%), Alexa Fluor 647 cadaverine, metal salts, acids, and bases were all purchased from Fisher Scientific. Thermal initiator 2,2’-Azobis[N-(2carboxyethyl)-2-methylpropionamidine]tetrahydrate (VA-057, 98%) was purchased from Wako as a solid and dissolved in deionised water to a concentration of 0.01 g/mL before use.

#### Mannose monomer synthesis

1,2,3,4,6-Penta-O-acetyl-α-D-mannopyranose (α or D) (5g, 0.0128 mols, 1 equivalent) was dissolved in 52.5 mL DCM and degassed with N_2_ for 15 minutes, then n-hydroxyethyl acrylamide added dropwise (2.56 mL, 0.0247 mols, 2 equivalents). After that, boron trifluoride diethyl etherate 0.6 mL/min (11.86 mL, 0.0961 mols, 7.5 equivalents) was added dropwise at a rate of 0.6 mL/min while keeping in an ice bath. The solution was left in ice for 1 hour, one hour later solution was kept at room temperature and left stirring for 36 hours. The reaction was monitored with TLC (100% Ethyl Acetate), which showed complete consumption of penta-acetyl mannose after 36 hours. The mixture was extracted using ice water twice, then the aqueous layer was extracted with DCM and the organic layers combined. These were washed with saturated NaHCO_3_ twice and once with brine, dried over sodium sulphate, then dry under vacuum to produce a pale yellow oil. This yellow oil was purified by flash column with hexane and ethyl acetate 20:80 eluent.

#### Mannose macro-RAFT agent synthesis

Mannose acrylamide monomer (731.5 mg, 2.48x10^−3^ mols, 20 equivalents) and DDMAT (50 mg, 1.24x10^−4^ mols, 1 equivalent) were dissolved in 5.593 mL MeOH, and then initiator, AIBN (407 μL, 2.48x10^−5^ mols, 0.2 equivalents, stock 10 mg/ml in methanol) was added. This solution was sealed and degassed with N_2_ for 10 minutes. This is then kept in oil bath at 75 °C for 3 hours stirring at 500 rpm. Reaction was checked by NMR in MeOD. Product was precipitated in diethyl ether, freeze for at least an hour to improve, then centrifuged and dried in vacuum oven.

#### Mannose macro-RAFT agent deprotection

Mannose macro-RAFT agent (400 mg, 3.8625x10^−5^ mols, 1 equivalent) was dissolved in 8 mL MeOH, at this point sample was taken for NMR (in MeOD). Then NaOMe (30% in MeOH, 41.2 μL, 3.8625x10^−4^ mols, 10 equivalents) was added followed by stirring for 1 hr at room temperature, and a pale a pale-yellow solid was formed. This was precipitated in diethyl ether, and dried in vacuum oven.

#### Emulsion Polymerisation

*α*D-mannose-CTA (40 mg, 0.0046 mmol, 1 equiv), was dissolved in 1.5 mL of distilled water. DPAEMA (110 *µ*L, 0.46 mmol, 100 equivs) or BMA (74 *µ*L, 0.46 mmol, 100 equivs) was added, along with VA-057 (55 *µ*L, 0.0013 mmol, 0.3 equiv). All reactions were stirred until dissolved, then the reaction was purged with nitrogen gas for 5 minutes. The reactions were immersed in an oil bath at 70°C for 90 minutes, stirred at 500 rpm. DLS: Z_av_ 100 – 200 nm, *PDI* 0.04 – 0.1.

For nanoparticles containing drug, rifampicin (5.43 mg, 0.0046 mmol, 1 equiv) was added in conjunction with the monomer, then reactions carried out as above.

The drug-loaded particles were then dialysed in distilled water for 24 hours before characterisation. DLS: Z_av_ 140 – 225 nm, *PDI* 0.05 – 0.17.

#### Drug Release

Nanoparticles were diluted to 20-fold in PBS and the pH adjusted using HCl to pH 7.4 or 4.5. Samples were taken at regular intervals, which were then spun at 14,000 rpm for 10 minutes to remove remaining nanoparticles. The supernatant was diluted 2-fold with MeOH then analysed via UV-Vis and HPLC for rifampicin content.

#### Alexa-Fluor 647 Conjugation

Nanoparticles were diluted to 10 mg/mL (2 mL, 1 equiv) in water. EDC (1 mg/mL in water, 0.3 equivs) and NHS (1 mg/mL in water, 0.3 equivs) were added to the solution, along with Alexa-Fluor 647 (1 mg/mL in water, 0.02 equivs). The reaction was sealed and left stirring at room temperature for 24 hours. The nanoparticles were then dialysed to remove any unconjugated dye before analysis.

#### NMR Spectroscopy

^1^H NMR and ^13^C NMR spectra were recorded on a Bruker Avance III HD 400 MHz spectrometer using deuterated solvent depending on the molecule to be measured, at a concentration of 40 mg/ml. For ^1H^ NMR spectroscopy the delay time was 2 s. Chemical shift values are reported in ppm downfield of a tetramethylsilane (TMS) standard. The results were analysed using Mestrenova software.

#### Transmission Electron Microscopy

Samples were prepared at 1 mg/mL in distilled water. 10 *µ*L was deposited onto a carbon-coated formvar copper grid, left for 1 minute, then wicked off. Samples were left to dry overnight before imaging on a JEOL 2100plus microscope with an accelerating voltage of 200 kV. Images were analysed using Image J software.

#### Dynamic Light Scattering (DLS)

Size measurements were taken on an Anton Paar Litesizer 500 at 25°C with a 40 mW single-frequency laser diode with a wavelength of 658 nm at a scattering angle of 175°. Nanoparticle samples were prepared at a concentration of 1 mg/mL in distilled water and measured in Omega cuvettes with an incubation time of 60 seconds at 25°C.

#### High-Performance Liquid Chromatography (HPLC)

HPLC measurements were carried out on an Agilient 1260 HPLC fitted with a fluorescence detector (FLD), which was set at 250 nm. Nanoparticles (125 *µ*L) were added to methanol (MeOH, 4.875 mL) to release the drug, then polymer was separated by centrifugation. The supernatant was removed and diluted by a factor of 2 in MeOH. Rifampicin standards were prepared in MeOH at concentrations of 50 – 25000 ng/mL All samples and standards were filtered through a 0.2 *µ*m poly(vinyl chloride) (PVC) filter. MeOH and pH 5 phosphate buffer were the mobile phase at a ratio of 70:30 for the first minute, then the methanol was increased up to 100% over the next 16 minutes, during which time the drug eluted. To restabilise the system the ratio was restored to 70:30 over 1 minute, then run at that ratio for 3 minutes before running the next sample. The flow rate was set at 0.5 mL/min for the entire experiment, and the injection volume was 20 *µ*L.

### Bacterial infection studies

#### Media and Reagents

Tween 80, glycerol, hygromycin B and RPMI rifampicin (97%) were purchased from Sigma Aldrich. Middlebrook 7H9 medium, Middlebrook 7H10 medium and oleic acid albumin dextrose-catalase (OADC) were purchased from Difco^TM.^; RPMI 1640 medium glutamine supplemented, Dulbecco’s modified Eagle’s medium (DMEM) with Glutamax and fetal bovine serum (FBS) were purchased from Gibco. DyLight 650 phalloidin (New England Biolabs), ProLong Gold antifade reagent with DAPI (Vector Laboratories), and LabTek II 4-well microscopy culture slides were purchased from Thermo Scientific. XTT Cell Proliferation Assay Kit (Generon Ltd), ACK red cell lysis buffer, accutase (Sigma-Aldrich) Company needed, accutase is a product), GM-CSF (Sigma), Pre-filled bead mill tubes (Fisher Catalog No.15-340-153).

#### Bacterial strains and culture conditions

*Mycobacterium tuberculosis*-H37Rv and BCG:eGFP (a kind gift from Maximiliano G. Gutierrez, The Francis Crick Institute) were grown at 37°C in Middlebrook 7H9 medium, supplemented with 10% v/v oleic acid albumin dextrose-catalase (OADC), 0.5% tween 80, and 0.02% v/v glycerol without or with 50 μg/ml hygromycin B. *Mtb* was handled in containment level (CL) 3 laboratory and BCG was handled in a CL2 laboratory.

On the day of the experiment, logarithmic (OD_600_ ∼ 0.5) cultures of *Mtb* and BCG were collected by centrifugation and the bacterial pellet was washed twice with phosphate-buffered saline (PBS) to remove residual culture media. Following the washes, the optical density of each bacterial suspension was measured using a spectrophotometer The cultures were then diluted to achieve the necessary bacterial density in the corresponding cell culture medium.

#### Human cell lines and culture conditions

Human promonocytic THP-1 cells (ATCC TIB-202) were cultured in RPMI-1640 medium with 2 mM L-glutamine (Gibco) and supplemented with 10% fetal bovine serum (FBS). A549 lung epithelial cells (ATCC CCL-185) were maintained in DMEM with Glutamax supplemented with 10% fetal bovine serum. Both cell lines were incubated at 37°C in a humidified atmosphere containing 5% CO₂.

#### Primary mouse-bone marrow derived macrophages (BMDMs)

Progenitor cells were isolated from the bone marrow of Balb/c mice (see below) and differentiated into macrophages following an established protocol with slight modifications^22^. Under aseptic conditions, the femur and tibia were harvested, and the surrounding tissue and muscle were carefully removed. The bones were washed in PBS, and both ends were trimmed with scissors. Bone marrow was flushed out using a 5-mL syringe filled with RPMI medium containing 10% FBS, attached to a 25-gauge needle. The extracted bone marrow was centrifuged at 200 g for 10 minutes at 4°C. The pellet was resuspended in 5 mL of RBC lysis buffer for 5 minutes at room temperature and centrifuged again at 200 g for 10 minutes. If needed, the lysis step was repeated. The cell pellet was then resuspended in RPMI medium with 10% FBS and passed through a 70-μm cell strainer. The cells were washed twice with the culture medium and counted using Trypan blue. One million cells were seeded into a petri dish containing 10 mL of culture medium supplemented with 20 ng/mL GM-CSF. On day 3, an additional 10 mL of GM-CSF-supplemented medium was added. On day 8, macrophages were harvested using accutase treatment for 15–20 minutes and seeded at the desired density for experiments.

#### Cytotoxicity assays

Cytotoxicity of RIF encapsulated nanoparticles against THP-1 macrophages and A549 epithelial cells were measured by XTT Cell Proliferation Kit. 1 x 10^4^ cells/well THP-1 cells were seeded in 96- well plate, maturation of THP-1 into macrophages was induced by incubation with 200 nM of phorbol 12-myristate 13-acetate (PMA) for 24 h. Similarly, 1 x 10^4^ A549 cells were seeded in 96-well plate 24 h prior to the assay. For cytotoxicity testing, cells were exposed to two-fold serial dilution of rifampicin encapsulated DPAEMA-pH-responsive mannosylated nanoparticles (RIF-DNP) or rifampicin encapsulated BMA-non-pH-responsive mannosylated nanoparticles (RIF-BNP) starting from 2 mg/ml to 0.0625 mg/ml. Some cells were exposed to RIF with a concentration equivalent to the RIF concentration encapsulated in nanoparticles. In control cells, an equal volume of sterile water was added. Cells were incubated with nanoparticles for further 24 hours; and then 50 µl of XTT working solution was added into each well. Cells were incubated for a further two hours with XTT solution and then absorbance of the samples was measured with a spectrophotometer at 450 nm and 650 nm. Percent viability of cells was measured using the following formula

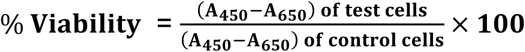

#### Uptake of nanoparticles in THP-1 cells

THP-1 cells were seeded at (5 x 10^5^ cells/well/ml) in 24-well plate or in a chamber slide, Maturation of THP-1 into macrophages was induced by incubation with 200 nM PMA for 24 h. After 24 h, cells were exposed to 100 μg/ml Alexa Flour 647-tagged DNP and BNP in cell culture medium. At each time point, nanoparticles were removed, cells were washed with PBS three times and fixed with 4% paraformaldehyde. Cells in the chamber slide were mounted with ProLong Gold antifade reagent containing DAPI, while cells in the plate were imaged soon after fixation. Images were taken at different time points starting from 30 minutes till 24 h with Cytation 5 (Agilent BioTek).

#### Efficacy of encapsulated RIF against Mtb and BCG in BMDMs

In this study, mouse bone marrow-derived macrophages were utilized to investigate the infection dynamics of *Mycobacterium tuberculosis*, *Mycobacterium bovis bacillus* Calmette-Guérin. BMDMs were seeded at a density of 2 x 10^5^ cells per well in a 24-well plate one day prior to infection.

For the infection, BMDMs were infected with either *Mtb* or BCG at a multiplicity of infection (MOI) of 10. The cells were incubated for 3 hours at 37°C in a humidified atmosphere with 5% CO_2_ to promote bacterial uptake through phagocytosis. After the incubation, extracellular bacteria were removed by thoroughly washing the macrophages with PBS.

Post-infection, the BMDMs were treated with either free rifampicin (RIF) or rifampicin formulations encapsulated in nanoparticles, specifically RIF-DNP (pH-responsive) and RIF-BNP (non-pH-responsive), at a RIF concentration of 1.00 µg/ml. The treated cells were incubated for either 1 or 3 days for *Mtb* and BCG, to evaluate the time-dependent effects of the treatments on bacterial viability. Control infected cells were left untreated to establish a baseline for bacterial growth and survival.

At each specified time point, the macrophages were lysed with cold water for 5 minutes to release intracellular bacteria. The lysates from *Mtb* and BCG-infected cells were serially diluted in PBS supplemented with Tween 80 and plated on 7H10 agar plates enriched with OADC (oleic acid, albumin, dextrose, and catalase) for colony-forming unit (CFU) enumeration. The plates were incubated at 37°C, and bacterial colonies were counted after 3 to 4 weeks for *Mtb* and BCG, allowing sufficient time for the slow-growing mycobacteria to form visible colonies.

#### Mice

In all experiments, we used 7–8-week-old female Balb/c mice were purchased from Charles River Laboratories, UK. All purchased animals were acclimatised in University of Warwick Rodent Facility for at least one week before starting any experiment. Mice were housed in individually ventilated cages, in a climate-controlled environment with a 12-hour light/12-hour dark cycle and food available ad libitum. Mice used for imaging were fed low auto-fluorescent diet (5V75, PicoLab® Verified 75 IF, Lab Diet), to avoid alfalfa related background in the imaging window starting one week before the experiment. All experiments were conducted under the licence PP9171369 and with local AWERB approval.

#### Seven-day repeat dosing and biodistribution of nanoparticles

To assess the toxicity of nanoparticles and monitor their distribution after 7-day repeat intravenous (i.v.) injection, mice were i.v. inject with with Alexa Fluor 647-DNP (100 µg/mouse in 50 µL PBS, n = 3), Alexa Fluor 647-BNP (100 µg/mouse in 50 µL PBS, n = 2), RIF (200 µg/mouse in 50 µL in PBS, n = 3) or PBS (50 µL, n = 2) only. Before the experiment, injected solutions were imaged as 10 µl drops in a petri dish to confirm comparable dye loading between DNPs and BNPs (Figure S4). All groups were imaged 1 h and 5 h, after the first two daily injections. From day 3 injection, only DPN and BNP groups were imaged once daily 1h after injection until day 7. 24 hours after the last injection, animals were imaged and culled.

Each day, all mice were weighed at the time of imaging and assessed for any signs of discomfort or toxicity using a standard inventory including grimace scale. For imaging, mice were anaesthetized with isoflurane and analysed using an *in vivo* imaging system (BioSpace Optima). Fluorescence was measured using the 637 nm excitation and 697 nm emission filter-set on 5s pictures. 24 hours after the last injection, all mice were culled, target organs were removed and assessed for fluorescence *ex vivo*.

#### Efficacy of free and encapsulated RIF in a BCG murine infection model

To evaluate the efficacy of rifampicin (RIF) encapsulated in mannosylated nanoparticles (RIF-DNP) in comparison to free RIF, mice were i.v. injected with 1 x 10^6^ *Mycobacterium bovis* Bacillus Calmette-Guérin (BCG) each in a volume of 50 µl in PBS (n = 20). One-week post-infection, four mice were euthanized to collect lung tissue, the primary target organ, as well as kidney and liver tissues as exemplar extrapulmonary sites. For the bacterial load assessment, all harvested organs were collected in pre-weighed tubes which were pre-filled with beads and weight of the tissue was determined. 1 ml of PBS-Tween (0.05%) was added in each tissue and then lysed with a homogeniser for 30 sec followed by 1 minute rest on ice. This was repeated for six cycles. The resulting lysates were serially diluted in PBS-Tween and plated on nutrient-rich 7H10 agar plates to enumerate colony-forming units (CFUs). On the same day, the remaining 16 infected mice were randomly divided into four groups and i.v. treated daily for one week with 50 µl of free RIF, RIF-DNP, free RIF and RIF-DNP, or PBS only as a control. For the single or combined treatment groups, each mouse was administered 6 µg RIF in total. On completion of the treatment course, all mice were sacrificed on day 8 and lung, kidney, and liver tissues were harvested and CFU was enumerated as above to assess the bacterial burden following the treatment.

## Results

### Preparation and characterization of RIF encapsulated mannosylated and pH-responsive polymeric nanoparticles

D-mannose-functionalized nanoparticles were synthesized via emulsion polymerization of hydrophobic monomers DPAEMA or BMA mediated by a RAFT macro-CTA, to yield nanoparticles with a pH responsive core based on poly(DPAEMA) (DNP) and non-pH responsive core based on poly(BMA) (BNP), respectively (**Figure S1**). Rifampicin (**Figure 1A**) was encapsulated by introducing the drug alongside the hydrophobic monomer. As polymerization progressed within the micelles and monomer droplets, the rifampicin was effectively sequestered within the hydrophobic core of the nanoparticles.

**Figure 1:**
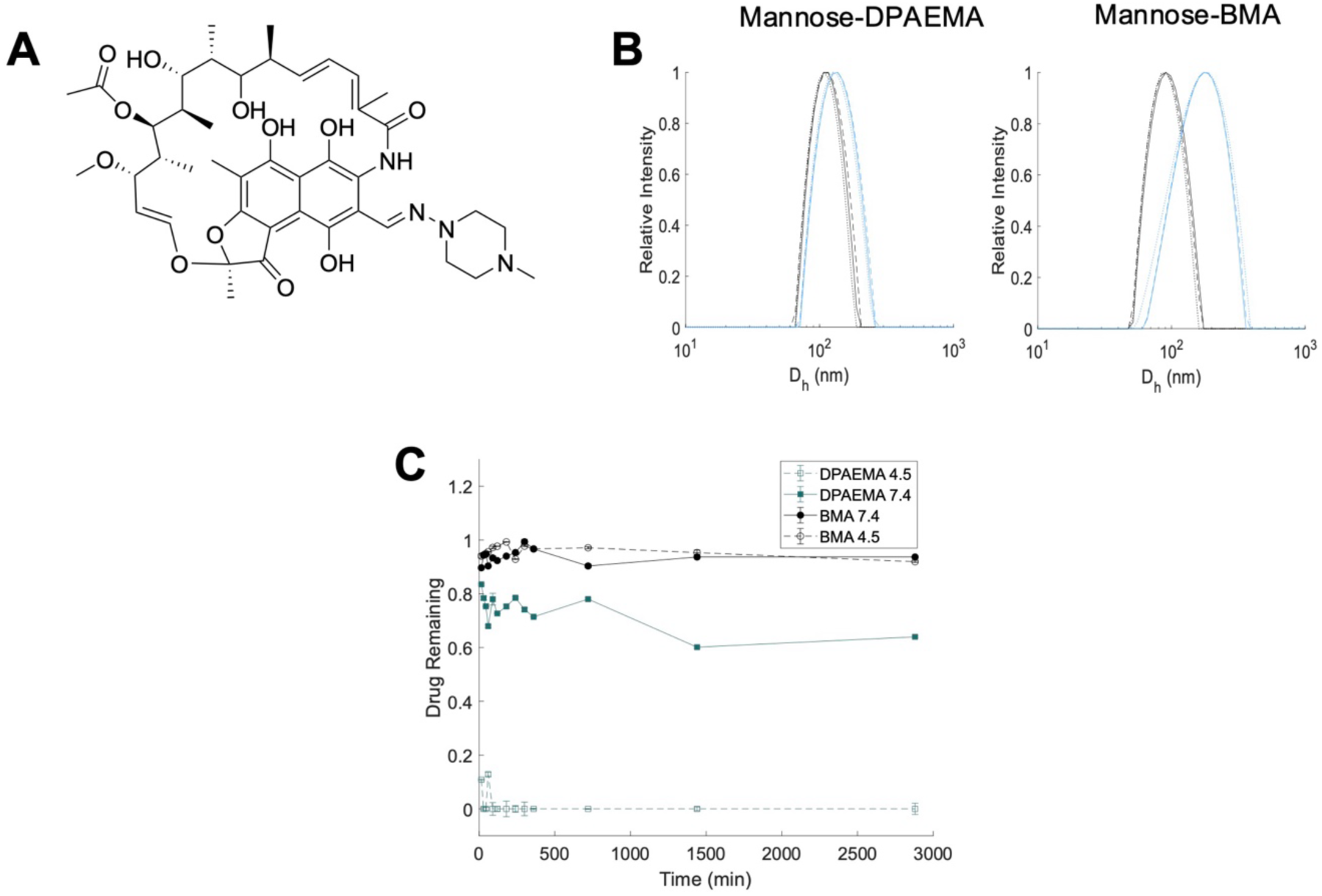
Rifampicin encapsulation and pH-responsive profile. (**A**) structure of rifampicin (RIF). (**B**) DLS of RIF encapsulated mannose nanoparticles before (black) and after (blue) dialysis. (**C**) shows the release of RIF from nanoparticles over time; Dashed lines represent a pH of 4.5, solid lines a pH of 7.4. N=3, error bars represent mean +/- standard deviation (SD).

Successful encapsulation was visually distinguished by the appearance of the purified product: the drug-loaded nanoparticles yielded an orange latex attributed to the intrinsic colour of rifampicin, in contrast to the characteristic white-blue latex of the drug-free controls. Following synthesis, the dispersions were dialyzed to remove non-encapsulated free drug. Dynamic Light Scattering (DLS) analysis confirmed the integrity of the particles post-dialysis, revealing a monodisperse size distribution (*PDI* 0.04 - 0.1) with hydrodynamic diameters ranging from 100 to 200 nm (**Figure 1B)**. Prior to dialysis, drug-loaded nanoparticles exhibited a reduced hydrodynamic diameter compared to their drug-free counterparts. This contraction is attributed to increased hydrophobicity within the core, leading to reduced water retention and tighter chain packing, alongside surficial stabilization provided by non-encapsulated rifampicin. Removal of free drug via dialysis led to an increase in particle size.

**Table 1** shows data for the rifampicin encapsulated nanoparticles. Size (Z_av_) and *PDI* were obtained from DLS and revealed DPAEMA nanoparticles (DNP) of 139 nm (*PDI* = 0.08) and BMA nanoparticles (BNP) of 176 nm (PDI = 0.15). To determine the concentration of encapsulated rifampicin, nanoparticles were disassembled in a solvent mixture of acetonitrile, water, and methanol. Following centrifugation to remove precipitated polymer, the supernatant was analysed against a standard calibration curve. HPLC analysis yielded a drug concentration of 0.25 mg/mL for both formulations, suggesting an Encapsulation Efficiencies (EE) of 18% for the BNP formulation and 17% for the DNP formulation. While these values fall within the broader literature range for similar synthesis methods,^23^ they are lower than the high efficiencies (50-90%) often reported for PLGA-based systems. ^23^

**Table 1.** Physical characterisation of nanoparticles based on poly(BMA) core (BNP) and poly(DMAEMA) core (DNP).

| Monomer | DP<sub>(th)</sub><sup>a</sup> | Z<sub>av</sub> (nm)<sup>b</sup> | PDI<sup>b</sup> | [Drug] (mg/mL)<sup>c</sup> | EE (%)<sup>d</sup> |
| --- | --- | --- | --- | --- | --- |
| BMA | 70 | 176 | 0.15 | 0.25 | 18 |
| DPAEMA | 70 | 139 | 0.08 | 0.25 | 17 |

<sup>a</sup> Degree of polymerisation; <sup>b</sup> determined by DLS; <sup>c</sup> Determined by HPLC; <sup>d</sup> Encapsulation Efficiency, determined by HPLC.

### Drug Release Analysis

To assess the suitability of the nanoparticles for macrophage targeting, drug release profiles were evaluated at physiological pH (7.4) and lysosomal pH (4.5). Nanoparticles were diluted 20-fold in PBS, pH-adjusted with 1 M HCl, and incubated over 48 hours. At scheduled intervals, aliquots were centrifuged to pellet intact nanoparticles. The supernatant was then diluted with methanol to precipitate any solubilized polymer chains, and the free drug concentration was quantified via UV-Vis spectroscopy (**Figure 1C**).

DNP particles exhibited a distinct pH-dependent release profile. At pH 7.4, where the polymer core remains unprotonated and hydrophobic, approximately 20% of the drug was released within the first 15 minutes, plateauing at 25% after 48 hours. This initial release is attributed to passive leakage driven by the concentration gradient upon dilution, likely facilitated by the moderate hydrophobicity of the DPAEMA core allowing for slight swelling and interfacial porosity. Conversely, at pH 4.5, the system demonstrated high efficiency, with 90% of the rifampicin payload released within 15 minutes, and 100% release within 1 hour. This rapid release was corroborated visually; the latex dispersion transitioned from translucent to transparent at pH 4.5, indicating complete core protonation and disassembly. In contrast, the control BNP particles showed no detectable drug release at either pH, confirming that the DPAEMA core is the sole driver of the pH-responsive mechanism.

These release characteristics are critical for addressing current challenges in TB therapy. Standard Fixed-Dose Combinations (FDCs) often suffer from reduced rifampicin bioavailability and patient non-compliance, factors that drive the emergence of drug-resistant strains. By protecting the cargo at physiological pH and ensuring rapid, triggered release within the lysosome, these nanoparticles offer a strategy to enhance bioavailability and mitigate degradation prior to reaching the infection site.

### Mannosylated DNP and BMA nanoparticles accumulate in macrophages

To facilitate intracellular tracking, nanoparticles were conjugated with the water-soluble fluorophore Alexa-Fluor 647 via EDC/NHS coupling (**Figure 2A**). This dye was selected for its emission wavelength (647 nm), which minimizes interference from biological auto-fluorescence in both *in vitro* and *in vivo* contexts. Successful conjugation was confirmed visually by the retention of the blue pigment within the latex dispersion and its absence in the dialysis water, with negligible effect on particle sizes (**Figure 2B**).

**Figure 2.**
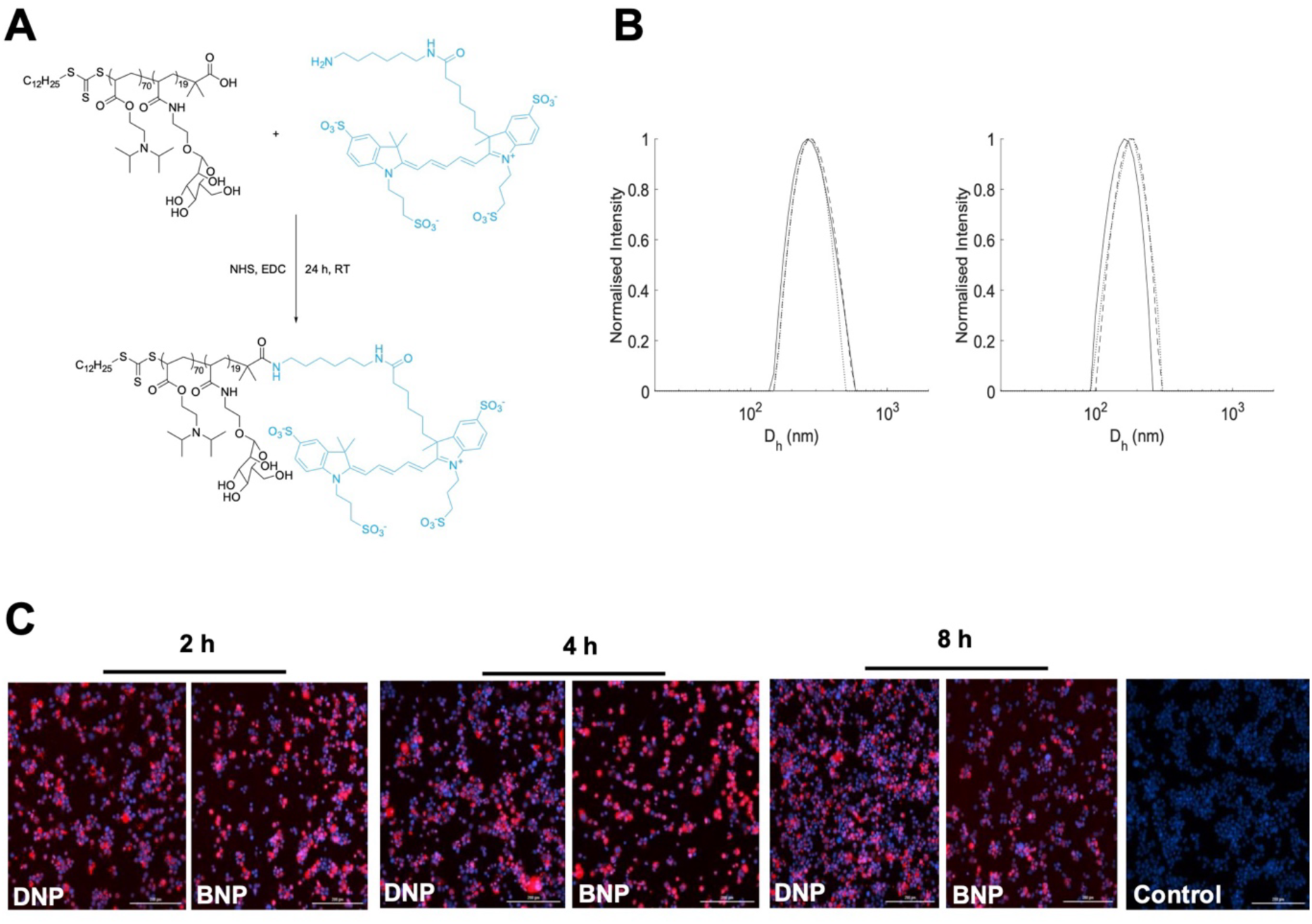
Fluorophore conjugation of nanoparticles and uptake into macrophages. (**A**) shows the reaction of conjugation of Alexa-Fluor 647 to mannose macro-RAFT agent via EDC and NHS coupling. (**B**) Dynamic light scattering (DLS) analysis of the nanoparticles conjugated with the Alexa dye. a) DPAEMA core, b) BMA core; size obtained from DLS is 190 nm and 175 nm respectively, with corresponding PDIs of 0.07 and 0.06. (**C**) To determine the uptake of nanoparticles, THP-1 cells were incubated with 100 μg/ml of nanoparticles. At indicated timepoints, cells were stained with DAPI and imaged. Red- Alexa Fluor 647 conjugated nanoparticle, Blue- DAPI stained nuclei, Scale bar-200 μm

Cellular internalization was evaluated in PMA-differentiated THP-1 macrophages. As hypothesized, no significant difference in uptake was observed between the BNP and DNP formulations (**Figure 2C**; **Figure S2**), attributable to their identical exterior mannose shell. However, both formulations demonstrated distinct time-dependent internalization kinetics, with fluorescence intensity increasing progressively over the incubation period. This aligns with previous observations contrasting mannose-functionalized particles against PEGylated controls, where specific receptor-mediated uptake drove higher, time-dependent accumulation^24^

### Nanoparticles are not toxic to human cells *in vitro*

To investigate the toxicity of the nanoparticles, THP-1 and A549 cells were exposed to various concentrations of RIF-DNP or RIF-BNP ranging from 2 mg/mL to 0.0625 mg/ml. Diluting nanoparticles in the same concentration range led to a change in intracellular RIF concentration, ranging from 13.5 - 0.4 µg/ml for DNP, 45.2 - 1.4 µg/ml for BNP. As a control, cells were treated with free RIF in equal concentration as that found in DNP. Viability of cells was measured by their ability to reduce a yellow tetrazolium salt to an orange formazan dye in a XTT assay. We did not observe any cellular toxicity at 24 hours for these particles and the free RIF at the concentrations tested **(Figure 3)**.

**Figure 3:**
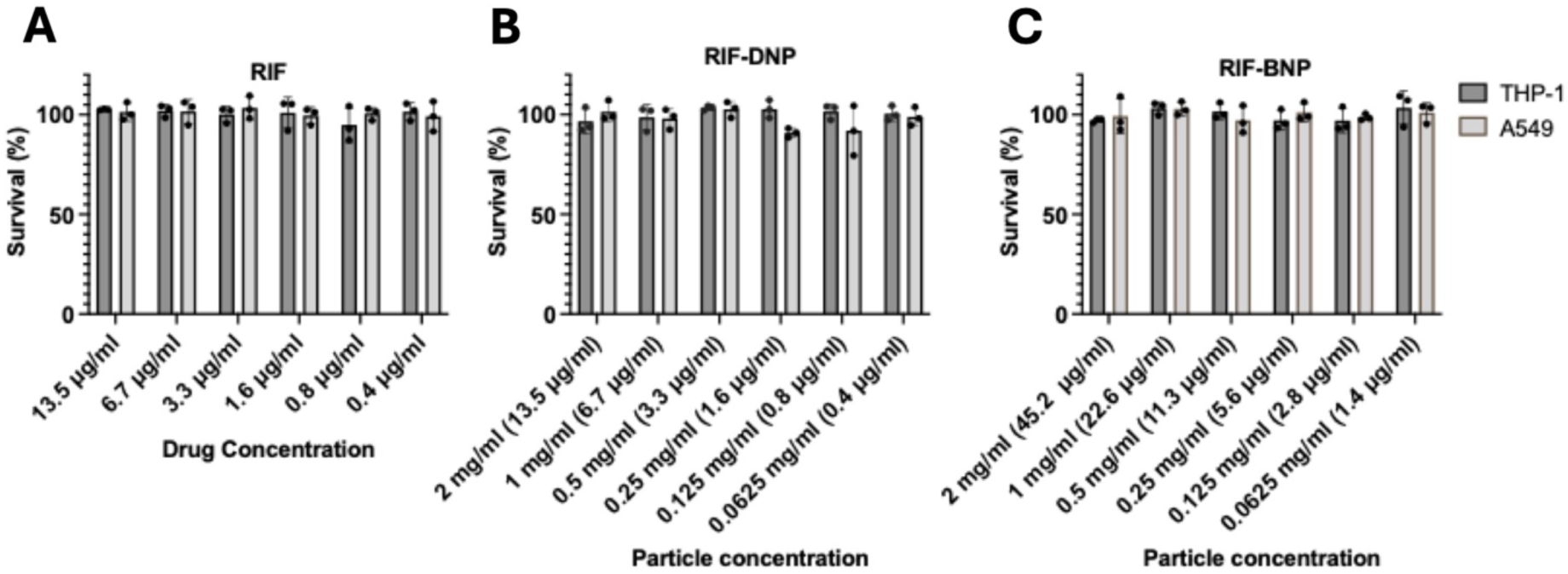
Nanoparticles are not cytotoxic. THP-1 macrophages and A549 epithelial cells were exposed to the indicated concentrations of drug-encapsulated nanoparticles or free drug for 24 h. Toxicity profiles of RIF alone (**A**), in non-responsive BMA core (**B**), and RIF encapsulated in pH responsive DPAEMA core (**C**) as measured by an XTT assay. Concentration of the encapsulated rifampicin in nanoparticles is shown in parenthesis.

### Rifampicin-encapsulated pH-responsive nanoparticles show superior efficacy over free drug against TB *in vitro*

To investigate the efficacy of encapsulated RIF, we first infected mouse-BMDM with *Mycobacterium bovis* BCG, a well-accepted model for TB. Three hours post infection, macrophages were washed with PBS to eliminate extracellular bacteria. Subsequently, the infected cells were exposed to RIF encapsulated in either pH-responsive nanoparticles (RIF-DNP) or non-pH-responsive nanoparticles (RIF-BNP), as well as free RIF. In all treatments, RIF was used at a standardized concentration of 1.00 µg/ml. After 24 hours and 72 hours post infection, cells were lysed for CFU enumeration. A significant bacterial reduction was observed when BCG-infected macrophages were treated with RIF-DNP when compared to RIF. Conversely, the bacterial load in cells exposed to RIF-BNP was comparable to the untreated control **(Figure 4A)**. This outcome aligns with the hypothesis that drug release does not occur in the non-responsive BMA core of RIF-BNP. RIF-DNP achieved a fold reduction in bacterial CFU of 4.92 ± 0.68 on day 1 and 48.77 ± 21.33 on day 3. In comparison, free RIF only demonstrated much lesser fold reductions of 1.77 ± 0.19 and 3.67 ± 1.23 on day 1 and 3 respectively. Since we used GFP-expressing BCG we also visualized the infected cells prior to lysis for quantifying bacterial counts. As depicted in **Figure 4B and Figure S3**, cells treated with RIF-DNP showed a significant reduction in bacterial load, whereas untreated cells and those treated with RIF-BNP exhibited high bacterial numbers

**Figure 4.**
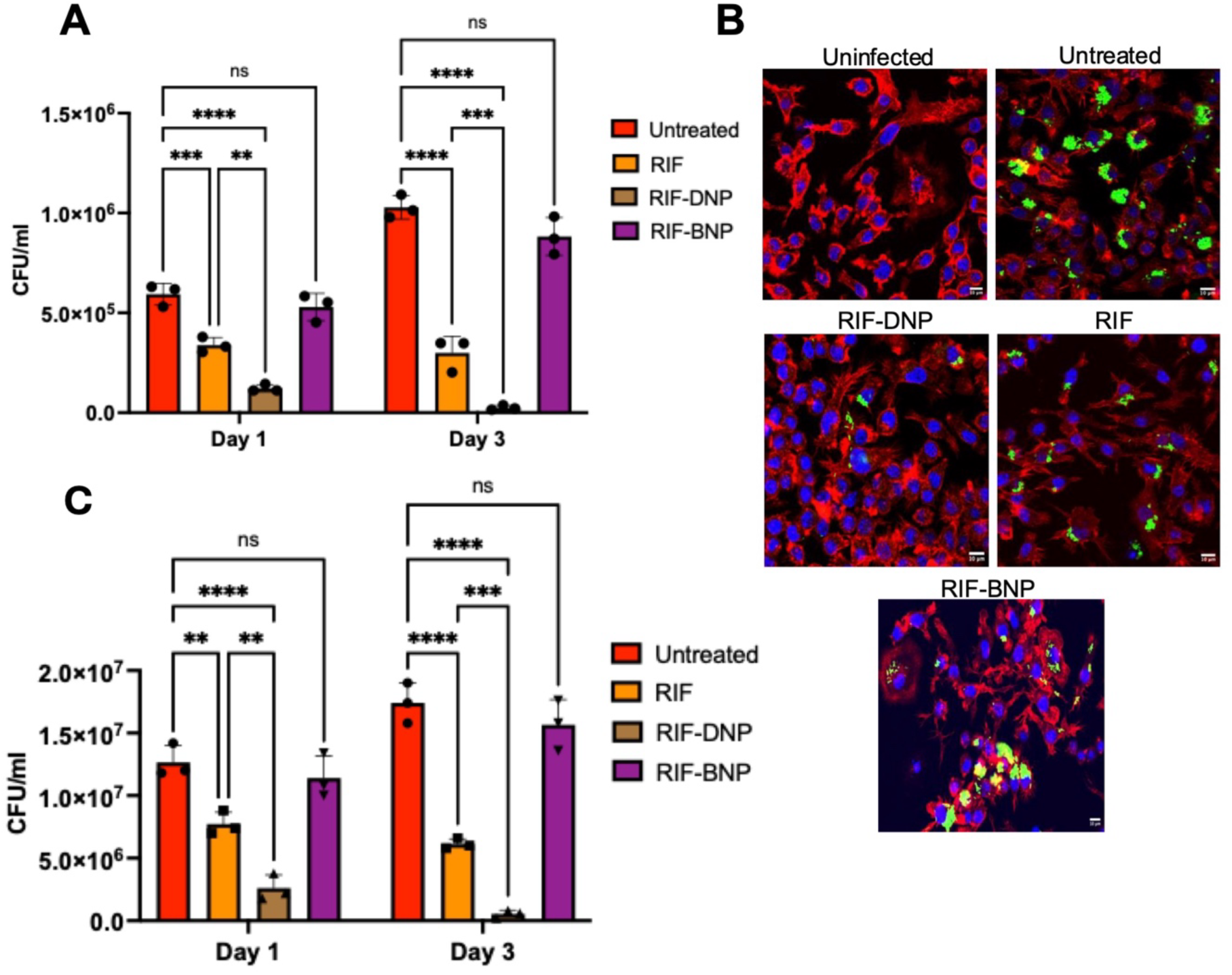
Encapsulated rifampicin shows better efficacy *in vitro*. After 24 h, mouse bone marrow derived macrophages (BMDM) were infected with BCG or Mtb for 3 hours at the MOI of 10, extracellular bacteria were removed by washing. Infected cells were exposed to free RIF or encapsulated nanoparticles (RIF-DNP or RIF BNP). On day 1 and 3 cells were lysed, and lysates were plated for CFU enumeration (**A**) colony counts for BCG. **(B**) BCG-infected macrophages were fixed, stained with phalloidin and intracellular bacteria were visualized by confocal microscopy green - BCG(GFP-tagged), red - actin (phalloidin-Alexa fluor-647), blue - nuclei (DAPI). Scale bar - 10 μm. (**C)** colony counts from in *Mtb*-infected BMDMs treated with free or encapsulated RIF. N = 3, mean +/- SD, \*\**P* < 0.01, \*\*\**P* < 0.001, \*\*\*\**P* < 0.0001 by an one-way ANOVA and Tukey’s multiple comparisons test.

Subsequently, we conducted a similar infection assay with the human pathogenic *Mtb* H37Rv strain. **(Figure 4C)**. RIF exhibited enhanced antibacterial efficacy when delivered through pH-responsive particles in *Mtb*-infected cells, with cells treated with RIF-BNP showing comparable bacterial numbers to the untreated control. A fold reduction in bacterial CFU of 5.375 ± 1.88 on day 1 and 45.9 ± 35.76 on day 3 was observed when treated with RIF-DNP, which was much higher than free RIF which showed a reduction of 1.65 ± 0.19 and 2.84 ± 0.18 on day 1 and 3, respectively.

Thus, our data show that rifampicin encapsulated in the pH-responsive nanoparticles (RIF-DNP) kill BCG and *Mtb* better than free rifampicin.

### Nanoparticles do not show toxicity *in vivo* after 7-day repeat dosing and do not accumulate in target organs of mice

To exclude toxicity and to study the biodistribution of DNPs and BNPs, the conjugated nanoparticles with the well-tolerated Alexa Fluor-647 were intravenously injected in mice, in parallel to injections of free RIF and PBS as controls, at the same time on 7 subsequent days. Fluorescence intensity was measured at 1 h and 5 h after i.v. injection on the first 2 days, and 1 h and 24 h after the last injection on day 7.

In line with the *in vitro* toxicity results, the weight of all mice was recorded daily throughout the study, revealing stable weight trends over time **(Figure 5D, Figure S4)**. Mice were also monitored daily for general appearance and any signs of discomfort, and none of the mice displayed indications for adverse health effects during the study.

**Figure 5.**
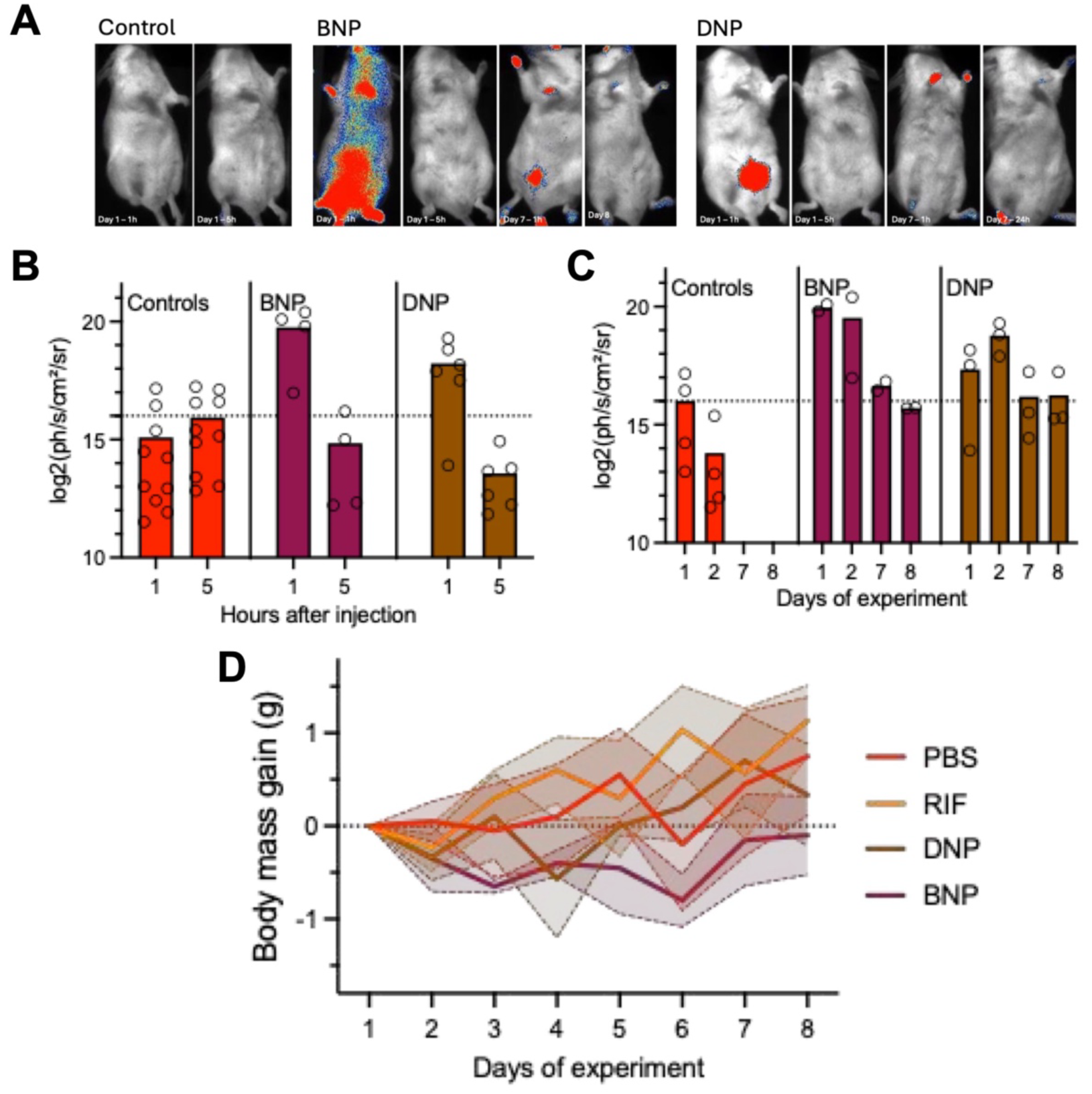
7-day nanoparticle injections in mice. Balb/c mice were i.v. injected daily with BNP (N=2) or DNP (N=3) nanoparticles (100 µg/mouse in 50µl PBS), or controls (free rifampicin (100 µg/mouse in 50 µl PBS (N=3) or 50 µl PBS (N=2)). After 1h and 5h, mice were anesthetized and imaged on day 1, 2. BNP and DNP were also imaged 1h and 24 h after last injection (day 8). (**A**) Representative imaging results and (**B**) quantitation of fluorescence intensity at 1h and 5 h after injection on day 1 or 2, and (**C**) at 1 h after injection on day 1, 2, 7 and 24h after injection (day 8). Of note: We pooled RIF and PBS treated groups for imaging quantification, as there was no detectable fluorescence in the imaging window of free drug, giving a better impression of the variation and background fluorescence. The dashed line in B & C indicates the level of background in the measurements. (**D**) Weight development of mice during 7-day repeated injections for all groups separately to also give an indication of the effect of RIF on the mice.

Notably, the nanoparticle groups exhibited significantly higher fluorescence than controls, i.e., free RIF or PBS, which displayed baseline signal counts **(Figure 5A, B, C)**. Among the nanoparticle formulations, BNP had a higher signal intensity than DNP. While both particle groups show a significant decrease in signal from 1 h to 5 h after injections a consistent trend to lower signal intensity was observed in the DNP group despite slightly a higher fluorescence for DNP when measured ex vivo (**Figure S5**). This decline suggests that the nanoparticles were progressively excreted or removed from circulation over time. Interestingly, the highest signal intensity was localized in the bladder region **(Figure 5A).**

### Rifampicin exhibits enhanced antimycobacterial efficacy *in vivo* when delivered via pH-responsive nanoparticles

We used an animal infection model to investigate the antibacterial effect of RIF encapsulated in mannosylated pH-responsive particles. Mice were infected intravenously with 1 x 10^6^ CFU/mouse. To assess bacterial load at the start of treatment, four mice were sacrificed one week after the initial infection, lungs, kidneys, and liver were carefully harvested and bacterial counts were determined. The lungs were evaluated for respiratory infection severity, as they are often the initial site of colonization, the kidneys and liver were included to assess the dissemination of the bacteria to other vital organs, which is a marker of infection progression. The lungs, kidneys, and liver harboured bacterial loads of log_10_ 5.083 ± 0.09, log_10_, 5.477 ± 0.10, and log_10_ 6.002 ± 0.10, respectively (**Figure 6A)** indicating that all three organs were colonized, with the liver showing the highest bacterial burden, in part due to the presence of highest number of macrophages (Kupffer cells), among all three organs.

**Figure 6.**
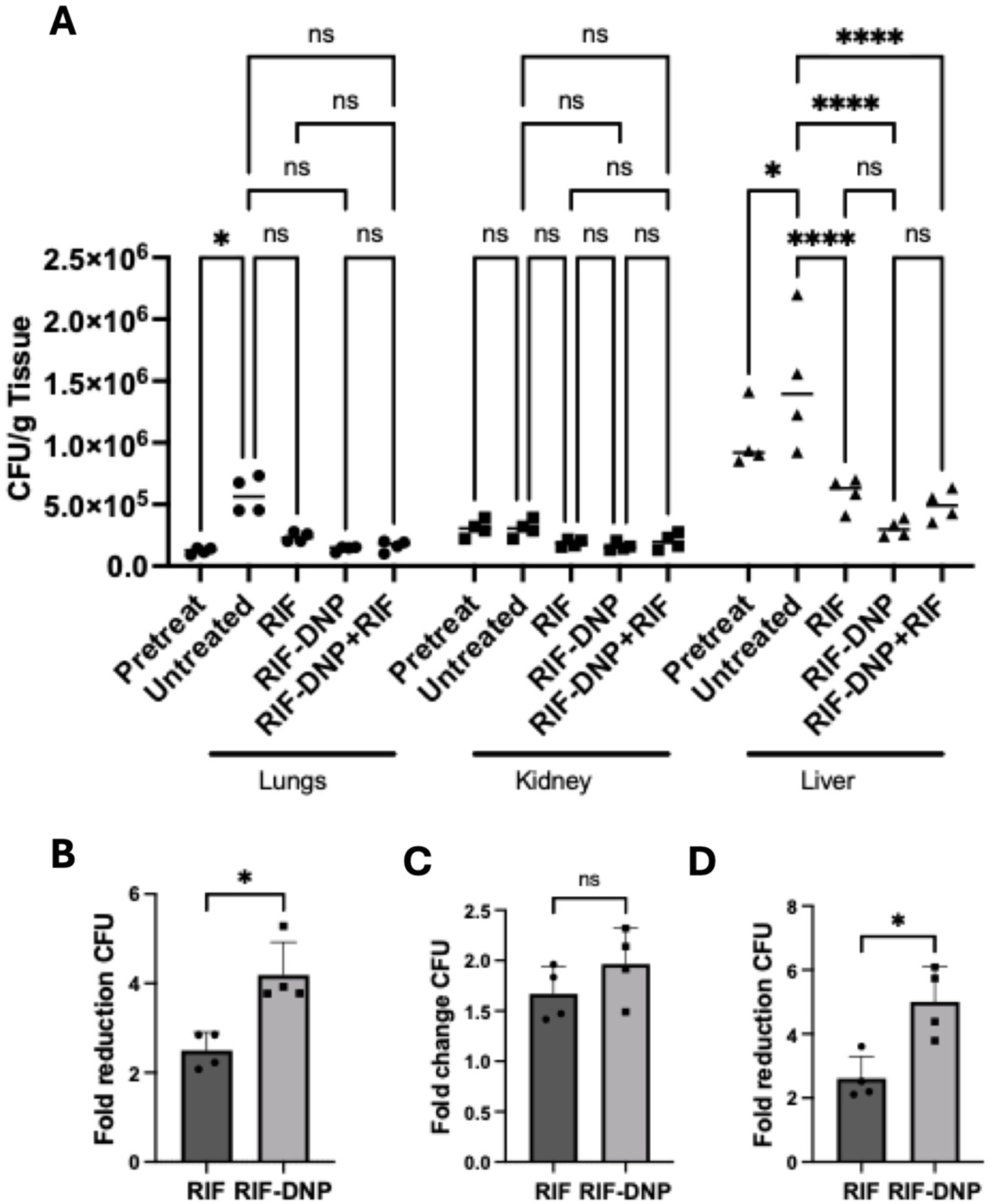
Efficacy of encapsulated rifampicin *in vivo*. Balb/c mice infected with BCG (1 × 10^6^ BCG/mouse). One week post infection, mice were treated with free RIF, encapsulated RIF (RIF-DNP-0.5 mg/kg) or encapsulated RIF + free RIF (RIF-DNP-RIF) daily. (**A**) CFU/g from lungs, kidney and liver obtained one week post-treatment are shown. (Pretreat = untreated group 1-week post-infection (prior to treatment), and untreated = 2-week post-infection untreated group). 2-way ANOVA analysis, with Tukey’s multiple comparison test,. ns- non-significant* P < 0.05, ****P < 0.0001. The fold reduction in CFU in free RIF- and RIF-DNP-treated groups relative to the untreated group in the lungs (**B**) kidney (**C**) and liver (**D**) are shown. N = 4/ group, mean +/SD *\* P < 0.05* as analysed by a Mann Whitney test.

After 1 week of infection, infected mice were treated intravenously with RIF-DNP, free RIF or RIF-DNP and free RIF daily for one week. We did not test RIF-BNP in this model because we did not observe activity of RIF encapsulated in these particles *in vitro* (Figure 4A). While both RIF and RIF-DNP reduced the bacterial loads compared to the untreated control (**Figure 6A**), RIF-DNP reduced the bacterial load in the lungs better than free RIF **(Figure 6A)**. Specifically, the bacterial reduction was 4.18 ± 0.73-fold for RIF-DNP but only 2.49 ± 0.4-fold for free RIF (**Figure 6B**). In contrast, no significant differences were observed in the kidneys between the two treatment groups **(Figure 6A)**, with bacterial reductions of 1.96 ± 0.35-fold for RIF-DNP and 1.67 ± 0.26-fold for free RIF (**Figure 6C**). In the liver, RIF-DNP treatment led to a significant reduction in bacterial burden compared to free RIF **(Figure 6A)**, with reductions of 5.0 ± 1-fold and 2.6 ± 0.68-fold, respectively **(Figure 6D)**.

The rationale for treating with RIF-DNP + free RIF was that in systemic infections, bacteria are known to reside both intracellularly and extracellularly; the intravenous infection model used in this study mimics a systemic infection. The combination therapy (RIF-DNP plus free RIF) resulted in bacterial reductions of 3.5 ± 1.34-fold, 1.52 ± 0.54-fold, and 3.0 ± 0.82-fold in the lungs, kidneys, and liver, respectively, thus showing a trend towards better efficacy than the RIF-DNP, although the differences between groups were not statistically significant (**Figure 6A**).

Thus, our data indicate that encapsulated rifampicin can treat BCG infections in mice more effectively than free rifampicin and combination therapy.

## Discussion

Treating tuberculosis (TB) effectively remains one of the top healthcare challenges worldwide. Novel delivery approaches are needed to enhance the efficacy of available anti-TB antibiotics as well as shorten the treatment time. Here we developed and evaluated mannosylated pH-responsive nanoparticles for targeted rifampicin delivery to macrophages, the key immune cells targeted by *Mycobacterium tuberculosis.* Our findings demonstrate that these nanoparticles effectively accumulate in macrophages, exhibit minimal cytotoxicity, and significantly enhance the intracellular antibacterial efficacy of rifampicin compared to both non-pH-responsive nanoparticles and free rifampicin. Furthermore, *in vivo* results indicate no overt toxicity at the used doses and drug accumulation in target organs and superior bacterial clearance in infected mice.

Nanomedicine has gained significant attention in modern healthcare due to its numerous advantages over conventional therapies, including optimized dosage regimens, reduced adverse effects, decreased drug degradation, lower dosing requirements, improved solubility and bioavailability, and consequently, enhanced patient compliance^25^. The successful uptake of our nanoparticles by macrophages aligns with previous studies that have shown mannose-functionalized nanoparticles achieve enhanced cellular internalization via the mannose receptor-mediated endocytosis pathway with mannose-modified nanoparticles demonstrating increased uptake compared to PEGylated counterparts^24,26,27^. We designed our nanoparticles to specifically release encapsulated drug in response to the acidic pH inside lysosomes, thereby preventing premature drug release. When macrophages were exposed to these, both pH-responsive and non-responsive formulations demonstrated significant uptake due to the presence of mannose on their surfaces. Notably, the uptake was time-dependent, supporting the premise that macrophages progressively internalize these nanoparticles for sustained drug delivery. In terms of cytotoxicity, neither the pH-responsive nor non-pH-responsive nanoparticles induced significant cell death in THP-1 or A549 cells even at the highest nanoparticle concentrations. These results reinforce the biocompatibility of our nanoparticle formulation, making them suitable for therapeutic applications.

Tuberculosis therapy poses unique challenges, as anti-TB drugs often exhibit poor efficacy against intracellular *Mtb*^28^. This limitation arises from factors such as inadequate cell membrane permeation, low intracellular drug accumulation, and non-specific target binding^5,28^. These challenges contribute to the development of drug-resistant strains and necessitate prolonged treatment regimens^29^. In recent years encapsulation of anti-TB drugs in pH-responsive nanoparticles have shown improved efficacy^30,18^. Poly(ethyleneimine)–poly(ethyleneglycol) (PEI–PEG) -coated isoniazid-carrying pH responsive nanoparticles were shown to kill TB *in vitro* and *in vivo* better than free isoniazid, although these were not specifically targeted to macrophages^30^. In another study mannose-modified lipid nanoparticles delivered isoniazid in a pH-sensitive manner, showing better efficacy than free drug^18^.

We encapsulated a key drug currently used in TB therapy, rifampicin (RIF), in an efficient and simple process. Our *in vitro* cell infection experiments provided compelling evidence of the superior antibacterial efficacy of pH-responsive nanoparticles in comparison to both non-responsive nanoparticles and free rifampicin. The significant bacterial reduction observed with RIF-DNP, but not for the non-pH-responsive BNP, suggesting that the pH-sensitive core effectively facilitates intracellular drug release within the lysosomal compartments of infected macrophages, thereby enhancing therapeutic efficacy. At acidic lysosomal pH, DPAEMA undergoes protonation causing the destabilisation of the polymer and, thus, releasing the drug. The BMA core lacks this pH sensitivity, retaining the drug within the nanoparticle even in lysosomal pH conditions. This targeted drug release aligns with the goal of achieving high intracellular drug concentrations while minimizing off-target effects. These findings are in line with previous studies that highlight the importance of pH-triggered drug release in maximizing intracellular antimicrobial effects against *Mycobacterium* species^24,31^ This targeted delivery significantly enhances the intracellular bioavailability of rifampicin.

Ensuring minimal toxicity and achieving good biodistribution of nanoparticles in organs are paramount factors for their successful application as vehicles for anti-tuberculosis drug delivery^32,18,30^. Our *in vivo* biodistribution data showed high uptake of nanoparticle formulations indicated by the fluorescence data. Importantly, the gradual decrease in fluorescence intensity of the nanoparticles over time suggests eventual nanoparticle clearance. While this should be further investigated at target organ level, this might indicate reduced risk of long-term toxicity. Crucially, no toxicity was observed in the mice in either the biodistribution nor efficacy experiments.

Our in vivo studies demonstrated that the efficacy of RIF-DNP was superior to free RIF a murine *Mycobacterium bovis* BCG infection model, which mimics many features of TB infection^33^. Our *in vitro* results suggest that encapsulated rifampicin penetrates intracellular compartments, where mycobacteria reside, more effectively, enhancing bactericidal activity. These findings provide a good rationale for developing this approach further. However, a weakness of our *in vivo* study is the use of a low dose of RIF (0.5 mg/kg) instead of recommended dose of 10 mg/kg^34^, which was necessary due to the low encapsulation efficiency of RIF (250 µg/mL). Further optimisation will allow to increase the loading efficiency of drugs in the particles and achieve a better therapeutic efficiency.

Given that intravenous BCG infection disseminates bacteria throughout the body, with both intracellular and extracellular populations, a combination therapy was also evaluated. Interestingly, the combination therapy of RIF-DNP with free RIF demonstrated an intermediate effect, likely due to the dual targeting of intracellular and extracellular bacteria. This observation highlights the potential benefits of combining nanoparticle-based drug delivery with conventional therapies to maximize treatment outcomes.

Overall, our findings provide strong evidence that mannosylated pH-responsive nanoparticles represent a promising strategy for enhancing rifampicin delivery and efficacy against tuberculosis. By improving intracellular drug delivery and demonstrating significant bacterial clearance, these nanoparticles have the potential to address key challenges associated with TB treatment, including drug resistance and suboptimal drug penetration. Future work should focus on optimizing the nanoparticle formulation, evaluating pharmacodynamics of these nanoparticles, and assessing their therapeutic potential in chronic and multidrug-resistant TB models to facilitate clinical translation.

## Acknowledgements

We are grateful for funding from the Medical Life Sciences Fund to PA and MU, a Monash Warwick Alliance fellowship to PA, a scholarship from the UKRI Warwick Centre for Doctoral Training in Analytical Science for HB and a Warwick-Wellcome Trust Translational Partnership Award for MU, RD and SP.

## Author contributions

**PA:** Formal analysis, Funding acquisition, Investigation, Methodology, Writing – original draft, Writing – review and editing

**HB:** Investigation, Methodology Writing – original draft

**RD:** Funding acquisition, Project administration, Resources, Supervision, Writing – review and editing

**SP:** Funding acquisition, Project administration, Resources, Supervision, Writing – review and editing

**MU**: Funding acquisition, Project administration, Resources, Supervision, Writing –, review and editing

