## Supplementary Information for "Targeting intracellular mycobacteria using novel antibiotic-loaded nanoparticles"

Meera Unnikrishnan*[1]

1. Warwick Medical School, University of Warwick, Coventry CV4 7AL, UK
2. Department of Chemistry, University of Warwick, Coventry CV4 7AL, UK
3. Faculty of Pharmacy and Pharmaceutical Sciences, Monash University, Parkville, VIC 3052, Australia

*Corresponding authors

Key words: tuberculosis, pH responsive, mannose, macrophages, nanoparticles, rifampicin

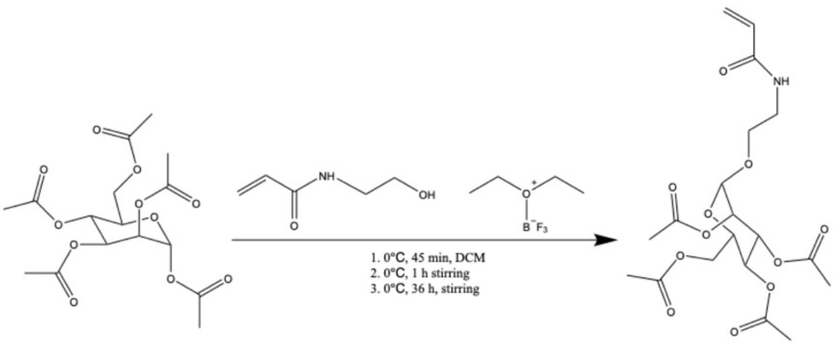


**Scheme 1.1:** Synthesis route of the mannose acrylamide monomer: Mannose pentaacetate is reacted with *N*-hydroxyethyl acrylamide (HEAA) in the presence of boron trifluoride ($BF_3$) until complete consumption of the pentaacetate is confirmed by thin-layer chromatography (TLC)

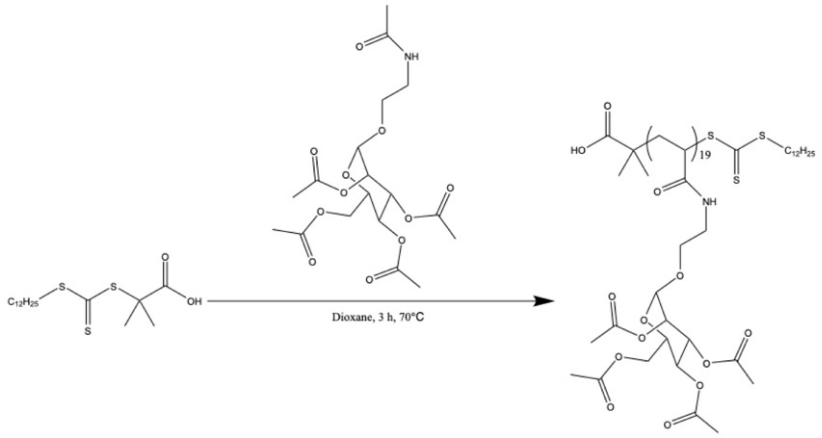


**Scheme 1.2:** Synthesis of mannose-CTA by RAFT solution polymerization

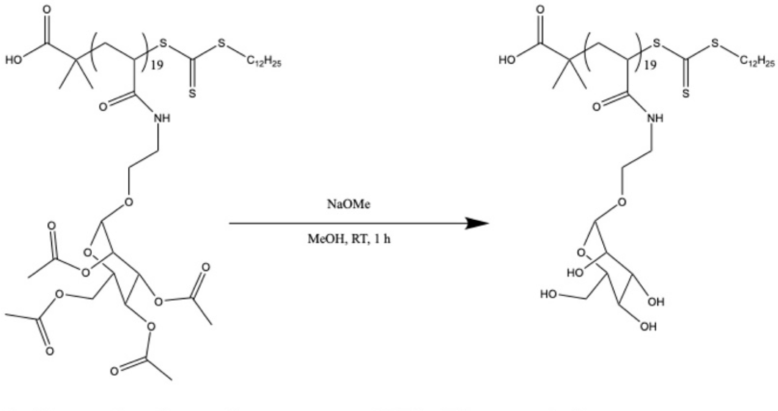


**Scheme 1.3:** Deprotection of mannose-CTA: The acetyl groups are removed using sodium methoxide (NaOMe), exposing the hydroxyl groups on the sugar, thereby enabling binding to the appropriate lectin receptor.

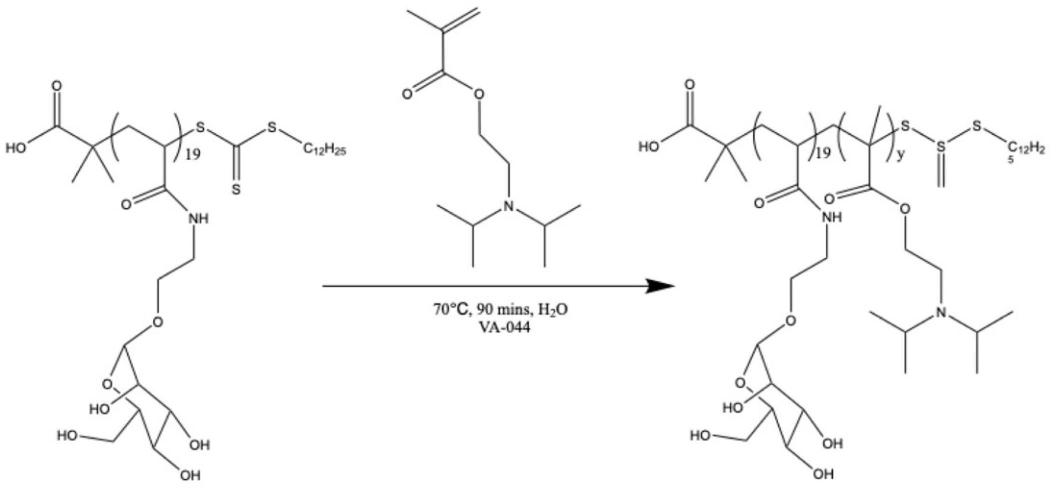


**Scheme 1.4:** Emulsion polymerization of mannose-CTA with DPAEMA

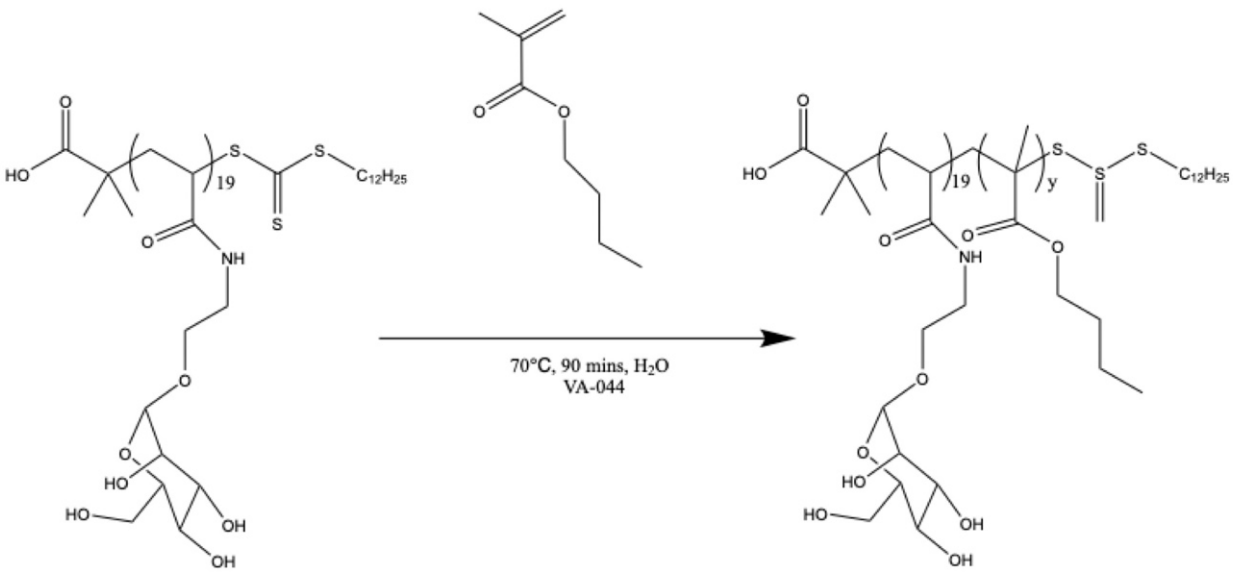


**Scheme 1.5:** Emulsion polymerization of mannose-CTA with BMA

**Figure S1. Schemes for nanoparticle synthesis**

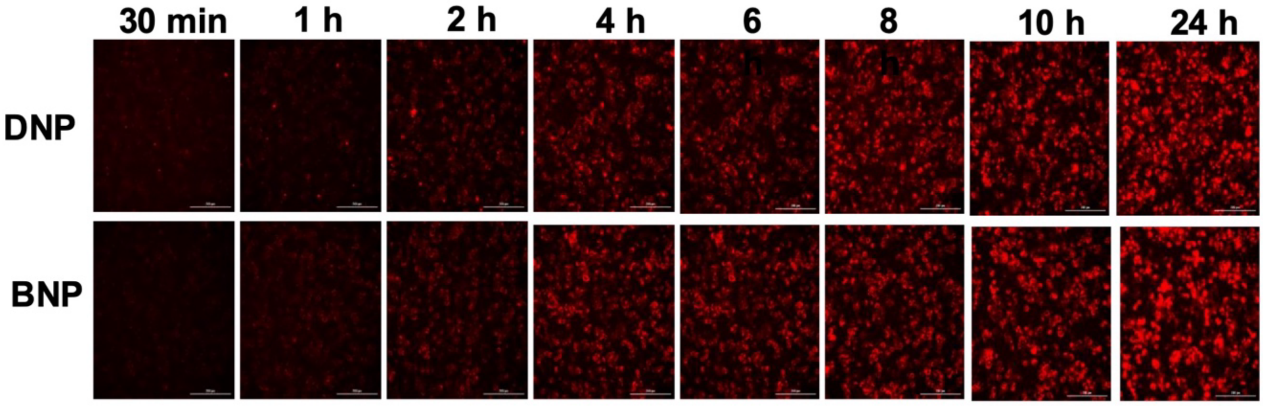


**Figure S2**. **Dynamics of nanoparticle uptake**. THP-1 cells were incubated with 100 µg/ml of nanoparticles, and at indicated time points, cells were fixed and imaged, red = Alexa Fluor 647 conjugated nanoparticle, Scale bar-200 µm

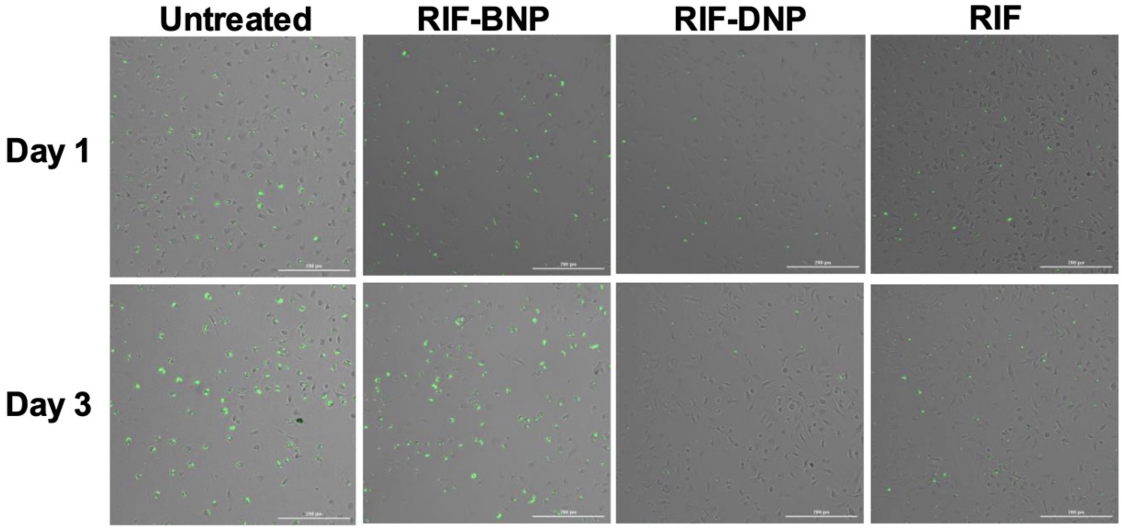


**Figure S3**. Mouse bone marrow derived macrophages (BMDM) were infected with BCG for 3 hours at the MOI of 10 and extracellular bacteria were removed by washing. Infected cells were exposed to drug encapsulated nanoparticles (RIF BNP or RIF DNP) and free drug (RIF). On day 1 and 3 cells were images just before cell lysis for CFU enumeration. BCG (green), cells (grey). Scale bar-200 μm

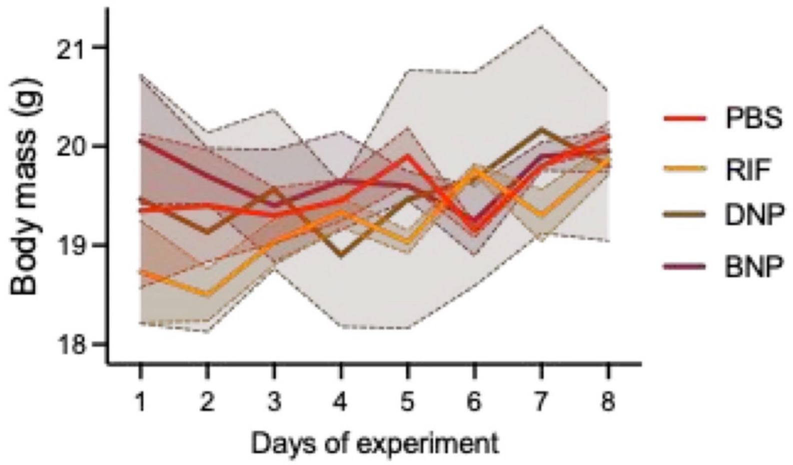


**Figure S4. Nanoparticles do not show negative effect on body mass development in 7-day repeat dosing in mice.** Mice were treated with PBS (N=2)), free drug (RIF, N=3), encapsulated rifampicin (RIF-DNP (N=3) or BNP (N=2)) and weighed each day. Mean and SD plotted for each group.

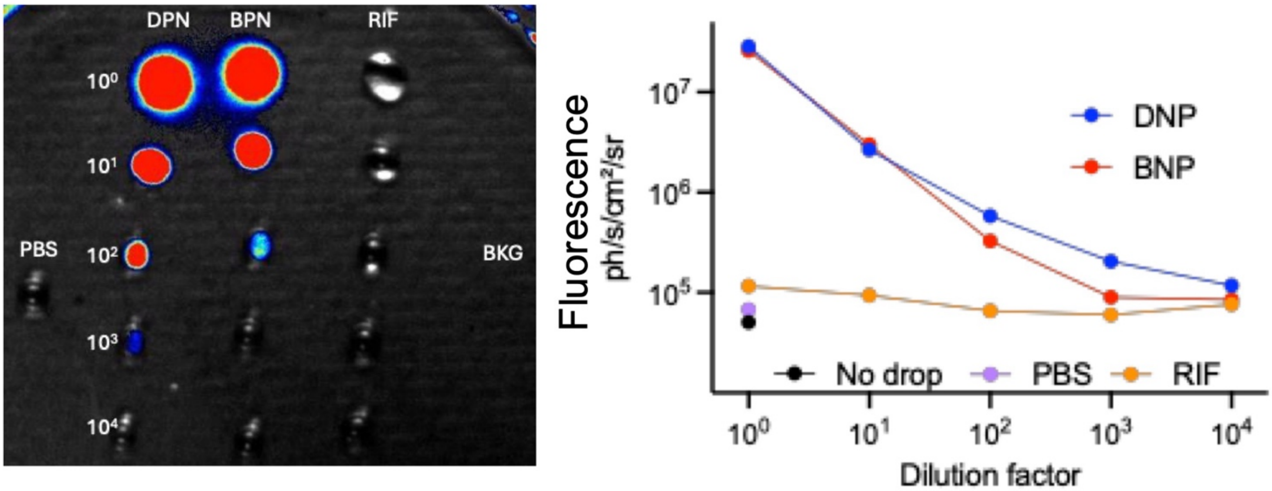


**Figure S5. Fluorescence values of dilution series of injected materials, control PBS and free drug imaged with the same setting as the mice and organs. Note the higher fluorescence values for DNP compared to BNP at 100-fold and high dilutions.**
